# The “neural shift” of sleep quality and cognitive ageing: A resting-state MEG study of transient neural dynamics

**DOI:** 10.1101/2021.07.23.453491

**Authors:** Roni Tibon, Cam-CAN, Kamen A. Tsvetanov

## Abstract

Sleep quality changes dramatically from young to old age, but its effects on brain dynamics and cognitive functions are not yet fully specified. We applied Hidden Markov Models (HMMs) to resting-state MEG data from a large cohort (N=564) of population-based adults (aged 18-88), in order to characterize transient neural networks and to relate their temporal dynamics to sleep quality and to cognitive performance. Using multivariate analyses of brain-sleep profiles and of brain-cognition profiles, we found that an age-related “neural shift”, expressed as decreased occurrence of “lower-order” brain networks, coupled with increased occurrence of “higher-order” networks, was associated with both increased sleep dysfunction and decreased fluid intelligence above and beyond age. These results suggest that poor sleep quality, as evident in ageing, may lead to a behavior-related shift in neural dynamics.

## 1 Introduction

With the increasing proportion of older adults in the worldwide population (Beard et al., 2016), there is a pressing need to understand the neurobiology of healthy ageing. Sleep quality changes dramatically from young to older age, and might be both causative and indicative of brain changes in healthy ageing (e.g., Feinberg et al., 1967; Fjell et al., 2018; Scullin & Bliwise, 2015). As people get older, sleep becomes more fragmented (e.g., Bliwise et al., 2009), less efficient (Gadie et al., 2017) and there is a decline in the quantity and quality of the “deep” stages of sleep, such as slow-wave sleep (SWS) and REM sleep (Ohayon et al., 2004; Varga et al., 2016). Among older adults, sleep problems have been associated with increased risk of developing cardiovascular disease (Wu et al., 2018), dementia (Shi et al., 2018), and mental health problems (Roberts et al., 2000). An outstanding question is how these variations in sleep quality are related to brain function and cognitive performance, which also change with age.

One way to investigate the relations between sleep quality and brain functioning is by relating sleep measures to resting-state functional connectivity. In recent decades, resting-state functional connectivity, measured mainly with functional magnetic resonance imaging (fMRI), have proved effective in distinguishing various patient groups from controls (e.g., Alzheimer disease, major depression, schizophrenia; see for example Lee et al., 2013). Substantial work has also used resting-state fMRI (rsfMRI) to examine the effects of age on functional connectivity (Andrews-Hanna et al., 2007; Chan et al., 2014; Ferreira & Busatto, 2013; Geerligs et al., 2015; Grady et al., 2016; Grady, 2008) with consequences for cognitive functioning (Bethlehem et al., 2020; Tsvetanov et al., 2016). Nevertheless, a recent attempt to relate resting-state functional connectivity to sleep quality, revealed no association between functional connectivity within or between resting-state networks and any objective or subjective sleep parameters (Lysen et al., 2020).

Decades-long investigation of sleep dysfunction and cognitive performance in healthy aging also yielded mixed results. Some studies suggest a link between poor sleep and reduced cognitive performance at older age (Feinberg et al., 1967; Hokett et al., 2021). For example, using a large cohorts of healthy older adults (N∼1500, 65+ years old), Tsapanou et al., (2017) showed that poor sleep quality and longer sleep duration were linked to low memory performance. In another large-scale multicohort study, sleep problems were associated with subjective cognitive decline in multiple cognitive domains including memory, naming, orientation and calculations (Tsapanou et al., 2019). Furthermore, it has been shown that, for older adults, subjective sleep problems in later life were predictive of cognitive decline as indicated by their score in the Mini Mental Status Examination (MMSE; Folstein et al., 1975). Nevertheless, others have indicated that these relations might be more limited. In particular, following a comprehensive review of seven correlational and experimental domains, Scullin & Bliwise (2015) concluded that in older adults, variability in sleep often does not relate to cognitive functioning, and that solely improving sleep may not reverse cognitive impairments.

The discrepancy between studies investigating the abovementioned associations between sleep quality, neural mechanisms, and cognitive performance in healthy ageing, might be due to several methodological factors. First, the relations between sleep quality and the brain, as measured with fMRI, might be affected by age-dependent confounding factors like cerebrovascular reactivity and head motion (Geerligs et al., 2017; Lehmann et al., 2017; Power et al., 2012; Tsvetanov et al., 2015). While some of these confounds, like neurovascular coupling, can be addressed by more sophisticated modelling (Tsvetanov, Henson, et al., 2021), others like head-motion are notoriously difficult to correct (Maknojia et al., 2019). Moreover, the brain-sleep relations might not be fully captured by static measures of functional connectivity. In particular, dynamic fluctuations in activity and connectivity can support flexible reorganization and coordination of neural networks (e.g., Allen et al., 2014), prompting the extension of functional connectivity to further include *dynamic* measures (e.g., Cabral et al., 2017; Vidaurre et al., 2018). Nevertheless, the fundamentally limited temporal resolution of fMRI (owing to the sluggish vascular response and relatively slow image acquisition times), precludes it from disclosing the potentially richer dynamics in brain connectivity above approximately 0.1 Hz. Finally, another potential source for this discrepancy might be the multivariate nature of the relations between sleep patterns, cognitive patterns, and the associated neural mechanisms. Indeed, a recent study in which age-related differences in sleep quality were associated with health outcomes across multiple domains, suggested that lifespan changes in sleep quality are multifaceted and not captured well by summary measures (Gadie et al., 2017).

To overcome these problems, in the current study we characterized transient resting-state neural dynamics that are related to sleep quality, cognitive performance and age using a multivariate approach. As an alternative to the indirect hemodynamic fMRI, we used magnetoencephalography (MEG) as a direct measure of neural activity sampled at 1kHz and higher. Unlike fMRI, MEG is not affected by age-related changes in vascular factors (Tsvetanov et al., 2015), and allows simpler and more robust methods for correcting head-motion artifacts (e.g., Taulu & Simola, 2006). We employed a recently developed analytical approach that offers the ability to measure dynamic functional connectivity in terms of transient states of stable patterns of activity and connectivity that last a few hundred milliseconds (Baker et al., 2014), well beyond the temporal resolution of fMRI. Furthermore, to account for the multifaceted nature of the factors considered in our study, we used Partial Least Squares (PLS) to relate patterns of neural dynamics to profiles of sleep quality and cognitive performance.

More specifically, we exploited a resting-state MEG dataset obtained from 564, population-based individuals, who were sampled uniformly across the adult-lifespan (18 to 88 years of age) as part of the Cam-CAN project (www.cam-can.org). In addition to MEG scanning, these individuals also completed a self-report sleep questionnaire (PSQI: Pittsburgh Sleep Quality Index; Buysse et al., 1989) and a wide range of cognitive tasks. We utilized the characteristics of the transient neural dynamics using Hidden Markov Models (HMM; Baker et al., 2014; Brookes et al., 2018; Hawkins et al., 2020; Vidaurre et al., 2016, 2017, 2018) as inferred in our previous study (Tibon et al., 2021). HMM is a data-driven method that identifies a sequence of “states”, where each state corresponds to a unique pattern of brain covariance that reoccurs at different points in time. By quantifying the time-series of MEG data as a sequence of transient states, the HMM provides information about the periods of time at which each state is active, enabling the characterization of its temporal dynamics. This technique had been used to identify neural dynamics in resting-state or task MEG datasets (Baker et al., 2014; Hawkins et al., 2020; Vidaurre et al., 2016), and was recently linked to age and cognition in a large-scale cohort (Tibon et al., 2021). The size of the Cam-CAN cohort allowed us to then take a multivariate approach, namely, to use PLS analysis to relate the temporal properties of the data-driven HMM states to profiles of sleep quality and cognitive performance.

Using similar methods, we have previously observed a “neural shift”, expressed as increased occurrence of brain states involving “higher-order” networks and decreased occurrence of brain states that involve early visual networks. This neural shift was associated with both increased age and decreased fluid intelligence, suggesting that it likely reflects reduction in neural efficiency rather than compensation (Tibon et al., 2021). For the current study, in accord with the inefficiency account, we predicted that the neural shift will be further associated with increased sleep dysfunction.

## 2 Materials and Methods

### 2.1 Participants

A population-based sample of 708 healthy human adults (359 women and 349 men) was recruited as part of Stage 2 of the Cambridge Centre Aging and Neuroscience (Cam-CAN; www.cam-can.org; (Shafto et al., 2014). Ethical approval for the study was obtained from the Cambridgeshire 2 (now East of England-Cambridge Central Research Ethics Committee), and participants gave full informed consent. Exclusion criteria included poor vision (below 20/50 on Snellen test; Snellen, 1862) and poor hearing (threshold 35 dB at 1000 Hz in both ears), ongoing or serious past drug abuse as assessed by the Drug Abuse Screening Test (DAST-20; Skinner, 1982), significant psychiatric disorder (e.g., schizophrenia, bipolar disorder, personality disorder), neurological disease (e.g., known stroke, epilepsy, traumatic brain injury), low score in the Mini Mental State Exam (MMSE; 24 or lower; Folstein et al., 1975), or poor English knowledge (non-native or non-bilingual English speakers); a detailed description of the exclusion criteria can be found in Shafto et al. (2014), Table 1. Of these, only participants who were considered for our previous study (Tibon et al., 2021; N=594, following the removal of 98 participants who did not have full neuroimaging data, 15 participants with poor MEG-MRI co-registration, and one participant who had no visits to one of the HMM states), and had full PSQI data (in all 7 measures of sleep quality), were considered for the current study. Thus, the final sample included 564 participants (age range 18-88).

### 2.2 Sleep measures

Sleep quality was assessed using the PSQI (Buysse et al., 1989), a well-validated self-report questionnaire designed to assist in the diagnosis of sleep disorders. The questions are grouped into seven components including overall sleep quality, sleep latency, sleep duration, sleep efficiency, sleep disturbance, sleep medication use, and daytime dysfunction due to sleepiness. Each of the sleep components yields a score on an ordinal scale, ranging from 0 (good sleep/no problems) to 3 (poor sleep/severe problems), with higher scores reflecting greater dysfunction. Scores were obtained from Gadie et al. (2017).

### 2.3 Cognitive tasks

Thirteen cognitive tasks, performed outside the scanner, were used to assess five broad cognitive domains, including executive function, memory, language, processing speed and emotional processing. The tasks are fully detailed in (Shafto et al., 2014). Task scores, also used in our previous study (Tibon et al., 2021; see Table 1 for full details), were obtained from Borgeest et al. (2018), in which missing data (<12% in all tasks) were interpolated using Full Information Maximum Likelihood (Enders & Bandalos, 2001) across the full Stage 2 sample (*n* = 708), as implemented in the Lavaan R package (Rosseel, 2012).

### 2.4 MEG resting-state data

MEG resting-state data included 4 temporal characteristics for 8 inferred brain states (i.e., 32 measures overall). The acquisition, preprocessing, and analysis pipeline are fully described in Tibon et al. (2021), and are summarized in Figure 1. In short, MEG resting-state data were recorded for 8 min and 40 s using a 306-channel VectorView MEG system (Elekta Neuromag, Helsinki), preprocessed using MaxFilter 2.2.12 software (Elekta Neuromag Oy, Helsinki, Finland), SPM12 (http://www.fil.ion.ucl.ac.uk/spm), and the OHBA Software Library (OSL; https://ohba-analysis.github.io/osl-docs/), and co-registered to each participant’s structural T1-weighted MRI. Source space activity was then estimated for each participant at every point of an 8 mm whole-brain grid using a single-shell lead-field model and a linearly constrained minimum variance (LCMV) scalar beamformer (Van Veen et al., 1997; Woolrich et al., 2011), parcelled into 38 regions of interest (ROIs; as in Colclough et al., 2015), and summarized by the first principal component across grid points within that parcel. The amplitude envelope of each parcel’s time-course was then calculated using a Hilbert transform.

**Figure 1.**
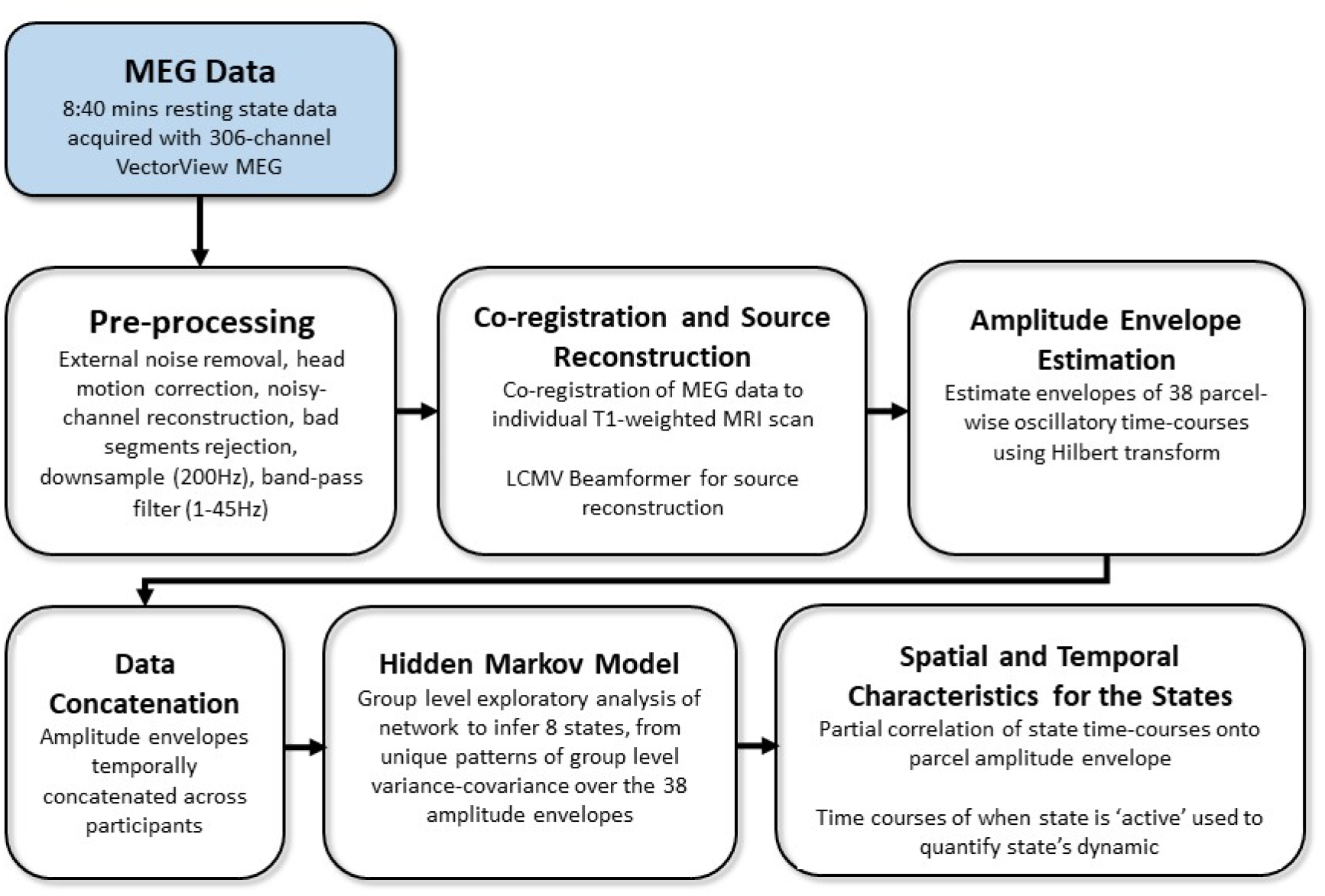
Overview of processing and analysis pipeline, adapted from Tibon et al. (2021).

Group-level exploratory analysis of networks (GLEAN; https://github.com/OHBA-analysis/GLEAN; Vidaurre et al., 2017) was then applied to the temporally-concatenated envelope data across all participants, in order to infer 8 brain states via Hidden Markov Modelling (HMM). HMMs describe the dynamics of neural activity as a sequence of transient events, each of which corresponds to a visit to a particular brain state. Each state describes the data as coming from a unique 38-dimensional multivariate normal distribution, defined by a covariance matrix and a mean vector. Therefore, each state corresponds to a unique pattern of amplitude envelope variance and covariance that reoccurs at different time points. The HMM state time-courses then define the points in time at which each state was “active” or “visited”. These estimated state time-courses, represented by a binary sequence showing the points in time when that state was most probable, were obtained using the Viterbi algorithm (Rezek & Roberts, 2005). Using these time-courses, the temporal characteristics of each state were quantified according to four measures of interest: (1) Fractional Occupancy (FO): the proportion of time the state was active; (2) Mean Life Time (MLT): the average time spent in the state before transitioning to another state; (3) Number of Occurrences (NO): the number of times the state was active; and (4) Mean Interval Length (MIL): the average duration between recurring visits to that state. The spatial and temporal characteristics of the HMM states are fully described in Tibon et al., 2021; Figures 2 and 3). The states include three distributed frontotemporoparietal networks (FTP1, FTP2, FTP3), a higher-order visual network (HOV), two early visual networks (EV1, EV2) and two sensorimotor networks (SM1, SM2). The spatial maps associated with the states are shown in Figure 2, Panel A.

**Figure 2.**
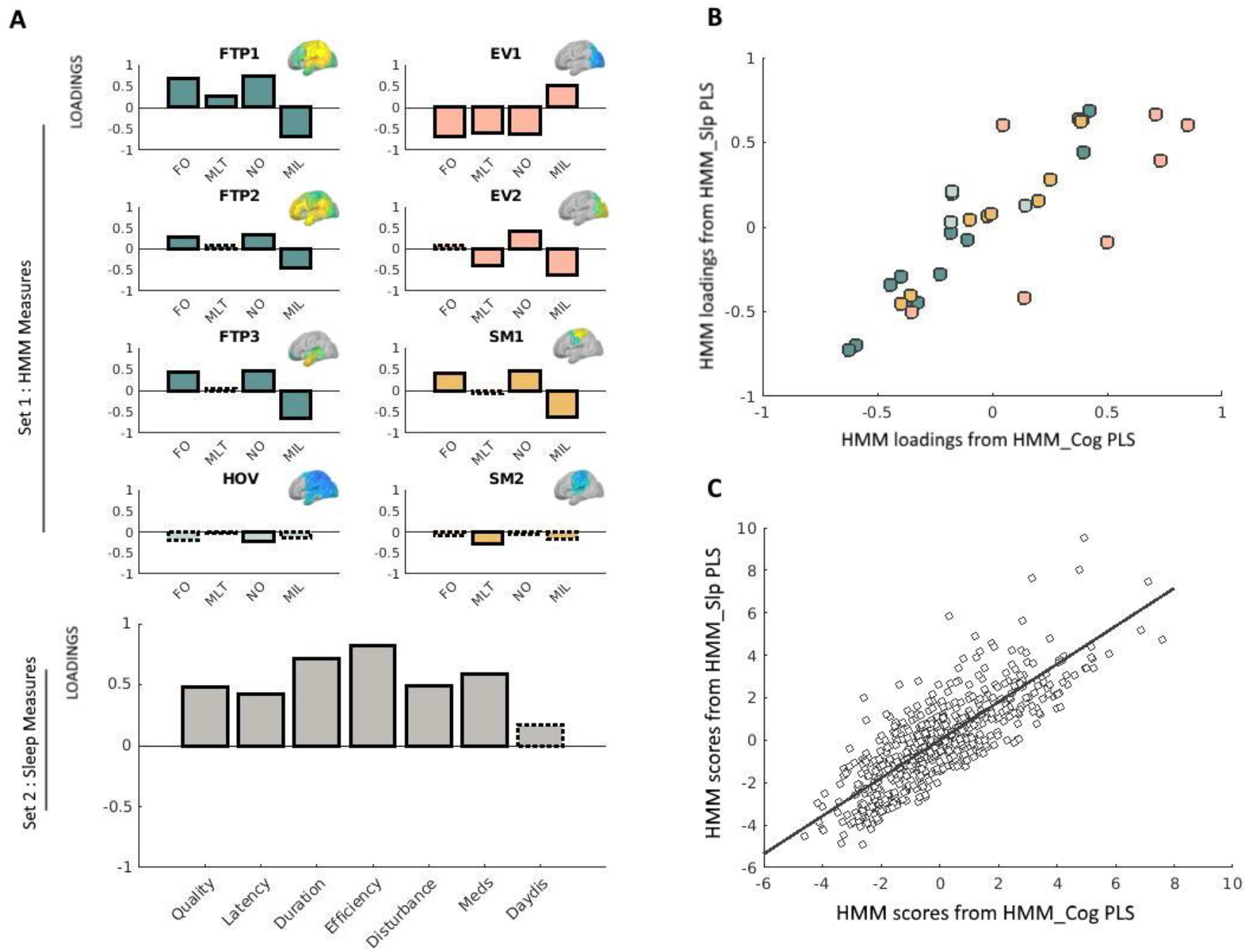
**(A)** Loadings obtained via the PLS analysis relating neural dynamics (HMM) measures with sleep measures. Solid outlines represent loadings greater than |.2|, whereas dashed outlines represent loadings smaller than |.2|. Loadings for network measures are shown in different colors, representing different types of states. HMM measures are indicated as *FO* (fractional occupancy, *MLT* (mean lifetime), *NO* (number of occurrences), and *MIL* (mean interval length). The various states are indicated as *FTP* (frontotemporoparietal), *HOV* (higher-order visual), *EV* (early-visual) and *SM* (sensorimotor). Corresponding HMM state maps (obtained from Tibon et al., 2021) are inset. For clarity, loadings for each network are shown separately, although in practice all equally contributed to a single PLS analysis. Loadings for the sleep measures are shown in gray (bottom-left panel). Sleep measures are sleep quality, latency, duration, efficiency, disturbance, sleep medication use (Meds), and daytime dysfunction (DayDis). **(B)** Scatter plot of the bivariate association between the loadings for the HMM measures obtained via the brain-sleep PLS analysis and the HMM measures obtained via the brain-cognition PLS analysis. Different colors reflect different types of states (FTP, HOV, EV, or SM), and correspond to the same color coding used in Panel A. **(C)** Scatter plot of the bivariate association between the scores for the HMM brain profile obtained via the brain-sleep PLS analysis and the HMM brain profile obtained via the brain-cognition PLS analysis. Two outliers were removed for the purpose of this visualization. This removal did not change the results (i.e., the correlation between these measures slightly increased and remained highly significant, *r* = .81, *p* < .0001). In this plot, each data-point represents one participant, whereas in Panel B above each data-point represents one measure.

### 2.5 Relating HMM states to sleep quality and to cognition (Partial Least Squares analysis)

For the brain-sleep analysis, we adopted a two-level procedure (Passamonti et al., 2019; Tsvetanov, Gazzina, et al., 2021). In the first-level analysis, we assessed the multidimensional relationships between temporal characteristics of the HMM states and sleep quality using PLS implementation in Matlab, the Mathworks Inc. This analysis describes the linear relationships between the two multivariate data sets by providing pairs of latent factors, as linear combinations of the original variables that are optimized to maximize their covariance. It is similar to the canonical correlation analysis (CCA) used in our previous study (Tibon et al., 2021), which instead maximizes the correlation between the latent variables. Although both CCA and PLS are useful to characterize relationships between two datasets, PLS has been suggested as a more appropriate tool for mixed datasets (Beaton et al., 2019; Grellmann et al., 2015), as is in our case, with the continuous and ordinal nature of the HMM and sleep data, respectively. All variables were z-scored before being subjected to the PLS analysis. First, we used a permutation-based PLS with 10,000 permutations to relate the 4 temporal characteristics across all 8 HMM states (Set 1, 32 variables) to the 7 sleep measures (Set 2).

Once we established the relationships between the HMM brain measures and the sleep measures, we asked whether the relationship between the HMM profile and the sleep profile (i.e., the relations between the participants’ scores on the latent variables obtained by the PLS analysis) varied with age, using a moderation analysis (see Tsvetanov et al., 2016, 2018, for a similar approach with different measures). Specifically, we constructed a second-level multiple linear model where HMM scores (on the latent variable), age, and their interaction term (HMM scores × age) were used as independent variables, and sleep scores (for the paired latent variable) were used as the dependent variable (all statistical tests were two-sided).

We then conducted another PLS analysis to relate the 32 HMM measures (Set 1) to the 13 cognitive measures (Set 2). This analysis resembles that in our previous study (Tibon et al., 2021) but uses PLS (instead of CCA) and a somewhat different sample (due to the exclusion of additional participants for which sleep data were not available). Similarly to our previous report, this analysis revealed a neural “shift” expressed as decreased occurrence of “lower-order” brain networks, and increased occurrence of “higher-order” networks, which was associated with decreased fluid intelligence. The reason to repeat this analysis in the current study was to obtain PLS scores and loadings that would be comparable with those obtained by the abovementioned PLS analysis, relating the HMM states to the sleep measures. Using these comparable scores, we asked whether the neural shift that was observed in our previous study, is also related to the pattern of sleep quality observed in the current study. To this end, we correlated both the HMM loadings and the profile scores (in two separate analyses), obtained by the first PLS analysis (i.e., which related HMM states to sleep quality) with the HMM loadings and profile scores obtained by the second PLS analysis (which related HMM states to cognition). Significant correlations would indicate that the pattern that was obtained for sleep quality and the pattern that was obtained for cognition are associated with the same neural pattern.

### 2.6 Data and code availability

Raw data from the Cam-CAN project are available from http://camcan-archive.mrc-cbu.cam.ac.uk, subject to conditions specified on that website. For a complete description of Cam-CAN data and pipelines, see Shafto et al. (2014) and Taylor et al. (2017). In addition, pre-processed mean data used for analyses and figures, together with the analysis code, are available on: https://osf.io/6nh5w/?view_only=ce2a24c365554a0ea49424900fa41a95.

## 3 Results

### 3.1 Relating HMM states to sleep measures

Our first step was to apply PLS to relate the 32 temporal characteristics of the HMM states (4 metrics for each of the 8 states) to the 7 sleep measures. This analysis identified one significant pair of latent factors (*p* = .01, based on a null distribution of 10,000 permutations). Figure 2, Panel A presents the loadings of this significant pair. For Set 1 (HMM data), the three frontotemporoparietal states (FTP1, FTP2, FTP3), and one of the sensorimotor states (SM1) showed positive loadings for the FO and NO measures, and negative loadings for the MIL measure. Furthermore, the first early visual state (EV1) showed negative loadings for FO, MLT, and NO and positive loadings for MIL. The second early visual state (EV2) displayed a similar pattern to that of EV1 for NO and for MIL, but not for NO (in which case the loadings for EV2 were positive). The higher order visual state (HOV) and the second sensorimotor state (SM2) were not strongly associated with this pattern of sleep quality (all loadings < .3). For Set 2 (sleep data), all of the components showed positive loadings (reflecting greater sleep dysfunction). The highest loadings were obtained for sleep efficiency, followed by sleep duration, and use of sleep medications. The lowest loadings were obtained for daytime dysfunction due to sleepiness. Taken together, poor sleep quality was associated with more and longer occurrences of states involving frontotemporoparietal regions and a state involving sensorimotor regions, and fewer, shorter occurrences of an early visual state.

The results obtained with the PLS analysis identified a neural shift that resembles the one related to cognitive decline as observed in our previous study (Tibon et al., 2021). However, before exploring this association further, we asked whether the relationship between the HMM brain profile and the sleep profile differs across age groups and exists beyond age. For this moderation analysis, we constructed a multiple linear regression model that included participants’ scores for the HMM profile, their age and the interaction (HMM profile × age) as predictors, and participants’ scores for the sleep profile as the dependent variable. The HMM scores were significantly associated with sleep scores after accounting for the main effect of age (*β* = .05, *t*(560) = 2.4, *p* = .017), demonstrating that the above brain-sleep relationship was not driven solely by age effects. The sleep scores were further associated with age (*β* = −.29, *t*(560) = −5.5, *p* < .001), verifying previous findings of age-related decline in sleep quality (e.g., Gadie et al., 2017). The interaction between age and HMM profile was not significant (*β* = .01, *t*(560) = .63, *p* = .5).

### 3.2 Relating HMM states to cognition (verification of previous findings)

Next, we applied PLS to relate the 32 temporal characteristics of the HMM states (4 metrics for each of the 8 states) to the 13 cognitive measures. This verifies the results of our previous study (Tibon et al., 2021), but uses a comparable method and sample to those that were used here, to test the association between HMM states and sleep. This analysis revealed a significant pair of latent variables (*p* = .01), depicting a pattern that highly resembled the pattern obtained in our previous study (two additional significant pairs depicted a different pattern, and were not explored further). The loadings obtained for this pair are shown in Supplementary Figure 1, together with the loadings obtained via CCA as in our previous study, in order to allow direct comparison. Although some of the loadings changed slightly, they were highly comparable with our previous results. Thus, here too, we observed that greater involvement of frontotemporoparietal states and reduced involvement of early visual states are associated with decreased fluid intelligence.

### 3.3 Correspondence between HMM profiles

Our final step was to investigate whether the neural shift that was associated with sleep dysfunction is the same as the one that was associated with decreased fluid intelligence. To this end, we correlated the HMM loadings obtained from the PLS analysis that related the HMM states to sleep measures, with the HMM loadings obtained from the PLS analysis that related the HMM states to cognitive measures. The correlation between these measures was highly significant, *r* = .83, *p* < .0001. However, as shown in Figure 2, Panel B, some states were more similar across the two PLS analyses than others. In particular, whereas the loadings obtained for the frontotemporoparietal, high-order visual, and somatosensory states were highly similar, noticeable variations were observed for the early-visual states. Finally, we correlated participants’ scores for the HMM profile obtained from the PLS analysis that related the HMM states to sleep measures with their scores for the HMM profile obtained from the PLS analysis that related the HMM states to cognitive measures. The correlation between these measures was also highly significant, *r* = .78, *p* < .0001.

## 4 Discussion

The results of our study show that transient neural dynamics, particularly those of frontotemporoparietal and early-visual states, are associated with sleep dysfunction. We found that increased sleep dysfunction is associated with increased occurrence of brain states involving “higher-order” networks and decreased occurrence of a brain state that involves an early visual network. Importantly, the same neural pattern was associated with decreased fluid intelligence and increased age (originally reported in Tibon et al., 2021, and verified in the current study). In our previous study (Tibon et al., 2021), borrowing from the approach that was applied to explain the posterior-to-anterior shift with ageing (PASA), commonly observed with fMRI during task (Davis et al., 2008; Glisky et al., 2001; Grady, 2012; Grady et al., 1994; Morcom & Henson, 2018; Nyberg et al., 2012; Park et al., 2004; Park & Reuter-Lorenz, 2009; Raz & Rodrigue, 2006; West, 2000), we considered two competing accounts for the shift from “lower” to “higher” networks that we have observed. The *functional compensation hypothesis* suggests that greater activation of higher-order regions serves to compensate for impairments in posterior brain regions, in order to maintain levels of cognitive performance. Alternatively, the *inefficiency account* suggests instead that the increased activation in higher-order regions reflects reduced neural efficiency or specificity. The crucial difference between these two accounts is that, whereas the functional compensation hypothesis predicts that the shift would correlate with better cognitive performance, the inefficiency account predicts the opposite pattern. Our previous findings that the neural shift was associated with worse cognitive performance, suggested that it represents reduced neural efficiency, thereby supporting the latter account. The current study provides further support for the neural inefficiency account by verifying the relations between the neural shift and decreased cognition (observed in our previous study) with another analytical approach, and by showing that the neural shift is further related to another maladaptive pattern—increased sleep dysfunction.

By using a novel data-driven method to infer brain states from MEG data, we were able to overcome some of the limitations of the more common use of fMRI to examine functional connectivity, such as confounding effects of vascular health, head motion and the ability to examine only very slow dynamics owing to low-frequency fluctuations of the fMRI response. Moreover, by using multivariate analyses to relate patterns of neural dynamics to patterns of sleep quality and cognitive performance, we were able to investigate multifaceted profiles that are not captured well by summary measures. Indeed, not all measures were equally associated with the neural pattern that we have observed. More specifically, as we observed in our previous study (Tibon et al., 2021), the cognitive measures of fluid intelligence were highly associated with the neural pattern, whereas those that measure crystallised intelligence were not. Similarly, a novel finding of the current study, was that while the observed neural pattern was associated with an overall increase in sleep dysfunction, the strength of the association with individual sleep measures varied: it was highly associated with sleep duration, efficiency, and sleep medication use (loadings > .5), moderately associated with sleep quality, latency, and disturbance (loadings > .2), and only loosely associated with daytime dysfunction due to sleepiness.

Using latent class analysis (LCA), Gadie et al. (2017) were able to classify individuals into four different classes representing “sleep types”, associated with distinct profiles of “sleep symptoms”. The probability of an individual showing the component profile associated with each class changed as a function of age for three of these classes labelled “good sleepers”, “inefficient sleepers”, and “delayed sleepers”. The component profile of the class labelled “inefficient sleepers” resembled the pattern observed in the current study (though notably, was also associated with increased sleep latency). This suggests that while age is associated with a general reduction in sleep quality, the neural shift observed in our study represents a more specific pattern of reduction. Importantly, our study shows that the relations between the neural shift and the sleep pattern cannot be fully attributed to age as a moderating factor (i.e., in our moderation analysis, these relations remained significant when controlling for age). Thus, in addition to its relation to ageing, the neural shift is further associated with inefficient sleep patterns that exist beyond age, throughout the entire adult lifespan.

An important factor that was associated with the neural shift observed in the current study was usage of sleep medications (3^rd^ highest loadings). Interestingly, studies have shown that some sleep medications are associated with changes in functional connectivity. For example, in a study by Pflanz et al. (2015), resting-state functional connectivity was investigated with fMRI, following seven-day diazepam administration. The authors found increased connectivity in response to diazepam administration in the medial visual network and middle/inferior temporal network. Furthermore, in a recent study (Frölich et al., 2020) with a sample of older adults (aged 55-73), infusion of midazolam resulted in increased rsfMRI functional connectivity between the dorsal default mode network and the posterior salience network. In the current study additional data regarding the kind of sleep medications and/or the dosage that was used are not readily available, nor can we infer any causal relations between sleep medications and functional connectivity. Therefore, potential associations between specific prescriptions and neural patterns cannot be explored further. It is important to note, however, that while it is unlikely that the neural shift is solely explained by usage of sleep medications (as other sleep measures also had substantial contribution), it is still associated with this factor to some extent. These relations between sleep medications and neural patterns should be explored further in future studies, especially given our finding that the neural pattern that is associated with increased use of medications is linked to reduced cognitive performance and to neural inefficiency.

In the current study, we observed a strong correlation between the loadings of the HMM measures obtained via the brain-sleep PLS analysis, and the HMM measures obtained via the brain-cognition PLS analysis, suggesting that they represent a highly similar pattern. Nevertheless, not all HMM measures were similarly comparable. In particular, while the HMM measures describing the frontotempoparietal states were highly similar, those describing the early-visual states were not. This is mainly because the observed sleep pattern was only reliably related to one of the early-visual states, whereas the observed pattern of cognitive performance was associated with both. We speculate that the reason for this discrepancy is that cognitive performance depend more on the visual system than sleep quality, and are therefore more strongly associated with particular neural patterns that involve these regions.

One limitation of our study is that we used cross-sectional data, which precludes direct inferences about ageing. However, we are not aware of any longitudinal MEG data on such a large, representative population, thus our results could be useful to generate hypotheses for future studies. The limitations of the MEG methods used in this study are thoroughly discussed in Tibon et al. (2021). In short, some properties of the assumptions and application of the HMM approach (e.g., group concatenation, Gaussian observation model, coarse parcellation of ROIs, and a-priori specification of 8 states) might result in oversimplification of the underlying neural dynamics. Nevertheless, some level of simplification is necessary for robust and interpretable modelling. Once the basic patterns are established, the parameters of the models can be adjusted to allow further optimization (see also discussion in Baker et al., 2014 and Quinn et al., 2018). Moreover, the current study used subjective reports of sleep quality. Although the PSQI questionnaire used for this purpose is well validated, it is yet to be determined whether the neural pattern that we have observed also correlates with objective measures of sleep quality (e.g., duration spent in Slow Wave Sleep; see review of various sleep measures in Hokett et al., 2021). Finally, the answers to the sleep questionnaire referred to the last month before the evaluation, and may not accurately represent long-term sleep patterns. Despite these limitations, our study verifies our previous findings, and further offers novel insights on the relationships between patterns of functional neural dynamics and sleep dysfunction in cognitive ageing.

## 5 Acknowledgments

The Cambridge Centre for Ageing and Neuroscience (Cam-CAN) research was supported by the Biotechnology and Biological Sciences Research Council (BB/H008217/1); R.T is supported by a British Academy Postdoctoral Fellowship (SUAI/028 RG94188); K.A.T is supported by the Guarantors of Brain (G101149); The funders had no role in study design, data collection and analysis, decision to publish, or preparation of the manuscript. We are grateful to the Cam-CAN respondents and their primary care teams in Cambridge for their participation in this study. We also thank colleagues at the MRC Cognition and Brain Sciences Unit MEG and MRI facilities for their assistance. The Cam-CAN corporate author consists of the project principal personnel: Lorraine K Tyler, Carol Brayne, Edward T Bullmore, Andrew C Calder, Rhodri Cusack, Tim Dalgleish, John Duncan, Richard N Henson, Fiona E Matthews, William D Marslen-Wilson, James B Rowe, Meredith A Shafto; Research Associates: Karen Campbell, Teresa Cheung, Simon Davis, Linda Geerligs, Rogier Kievit, Anna McCarrey, Abdur Mustafa, Darren Price, David Samu, Jason R Taylor, Matthias Treder, Kamen A Tsvetanov, Janna van Belle, Nitin Williams; Research Assistants: Lauren Bates, Tina Emery, Sharon Erzinçlioglu, Andrew Gadie, Sofia Gerbase, Stanimira Georgieva, Claire Hanley, Beth Parkin, David Troy; Research Interviewers: Jodie Allen, Gillian Amery, Liana Amunts, Anne Barcroft, Amanda Castle, Cheryl Dias, Jonathan Dowrick, Melissa Fair, Hayley Fisher, Anna Goulding, Adarsh Grewal, Geoff Hale, Andrew Hilton, Frances Johnson, Patricia Johnston, Thea Kavanagh-Williamson, Magdalena Kwasniewska, Alison McMinn, Kim Norman, Jessica Penrose, Fiona Roby, Diane Rowland, John Sargeant, Maggie Squire, Beth Stevens, Aldabra Stoddart, Cheryl Stone, Tracy Thompson, Ozlem Yazlik; and administrative staff: Dan Barnes, Marie Dixon, Jaya Hillman, Joanne Mitchell, Laura Villis. The authors declare no competing interests.

## Supplementary Materials

**Supplementary Figure 1.**
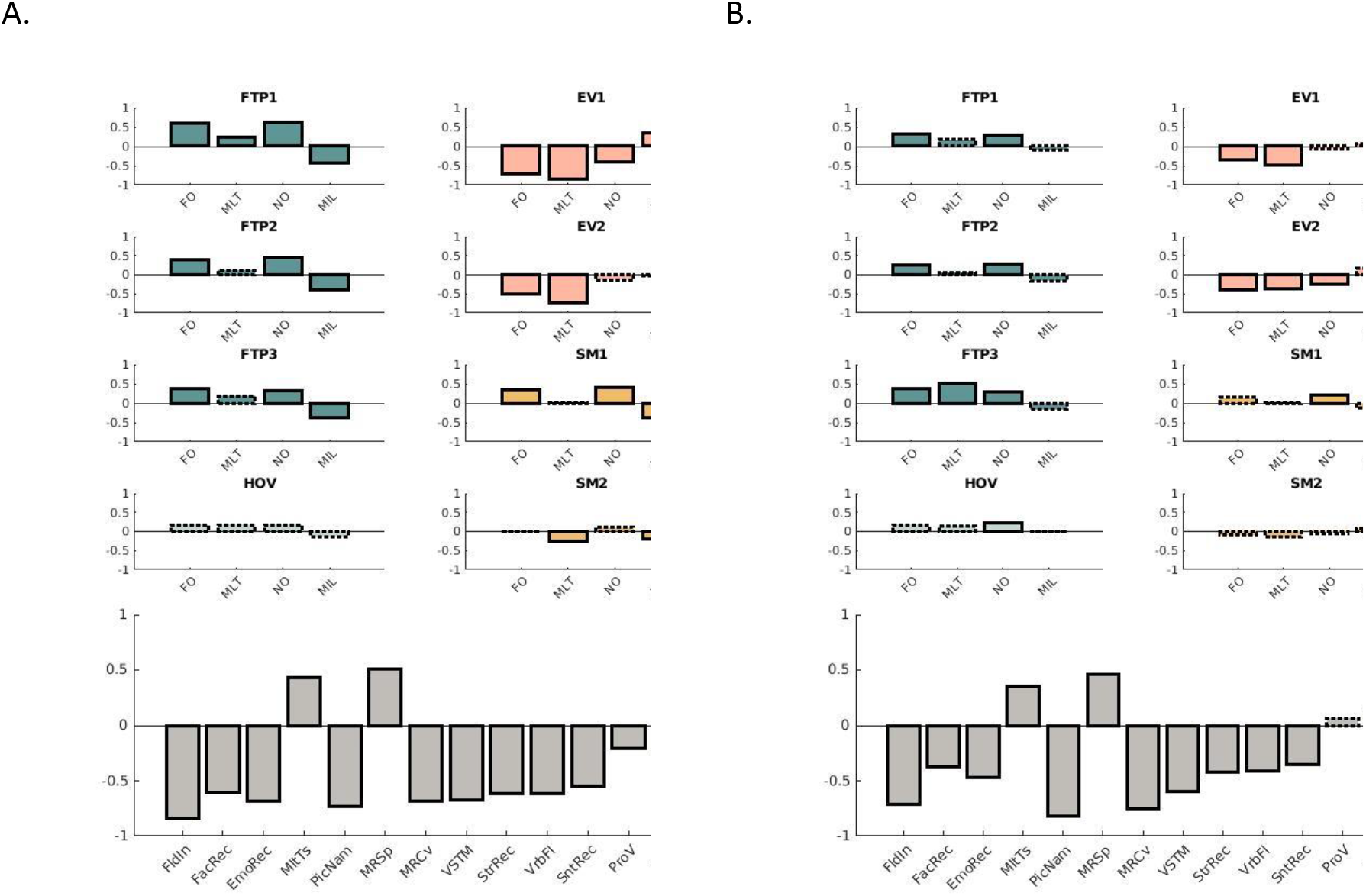
Loadings obtained via the analysis relating brain network (HMM) measures with cognitive measures using PLS (Panel A) and CCA (Panel B). Overall, the results are highly similar. The CCA results displayed in Panel B were obtained using the same method reported in Tibon et al. (2021), but on a subset of the sample (with N=564 instead of N=594) due to the exclusion of participants with no sleep measures.

